# The impact of molecular variants, crystallization conditions and space group on structure-ligand complexes: A case study on Bacterial Phosphotriesterase Variants and complexes

**DOI:** 10.1101/2023.05.31.542999

**Authors:** Orly Dym, Nidhi Aggawal, Yaacov Ashani, Shira Albeck, Tamar Unger, Shelly Hamer Rogotner, Israel Silman, Dan S. Tawfik, Joel L. Sussman

**Author notes:** **Synopsis** This study provides valuable insights into the challenges and considerations involved in studying the 3D structure of proteins with bound ligands, highlighting the importance of careful experimental design and rigorous data analysis to ensure the accuracy and reliability of the resulting protein-ligand structures.

## Abstract

While attempting to study the 3D structure of proteins with bound ligands, one often encounters considerable difficulties. We illustrate, as an example, the bacterial enzyme phosphotriesterase and specifically examine the effects of multiple factors such as the molecular constructs, ligands used during protein expression and purification, crystallization precipitance, and space group on the visualization of molecular complexes of organophosphate ligands bound to the enzyme.

We analyzed twelve crystal structures of the different phosphotriesterase constructs derived by directed evolution in both apo and holo forms (in complex with organophosphate analogs), with resolutions up to 1.38 Å. Crystals obtained from three different crystallization conditions, crystallized in four space groups, with and without N-terminal tags, were utilized to investigate the impact of these factors on visualizing molecular complexes of ligands bound to the enzyme. The study revealed that residual tags used for protein expression can lodge in the active site and hinder ligand binding. Additionally, the space groups in which the proteins are crystallized can significantly impact the visualization of the organophosphate ligands bound to the phosphotriesterase. The study also reveals that the crystallization precipitants can compete with and even preclude ligand binding, leading to false positives or the incorrect identification of lead drug candidates, which is particularly crucial for ligands with pharmacological and toxicological contexts.

Overall, this study provides valuable insights into the challenges and considerations involved in studying the 3D structure of proteins with bound ligands, highlighting the importance of careful experimental design and rigorous data analysis to ensure the accuracy and reliability of the resulting protein-ligand structures.

## 1. Introduction

Structure-based drug design is a powerful tool in drug discovery, allowing researchers to predict how a potential drug will interact with its target protein. X-ray crystallography is a widely used method to obtain detailed information about the protein-ligand complex, but it also has some limitations that can affect the accuracy and reliability of the obtained protein-ligand structure. About 75% of the ∼175,000 X-ray crystallographic studies of biomacromolecules in the Protein Data Bank (PDB) (Berman *et al*., 2000; Sussman *et al*., 1998) contain at least one of nearly 5,000 unique ligands. Some ligands may be present unintentionally due to the purification or crystallization process (Dym *et al*., 2016), while others are deliberately added to the sample to study protein function or as part of structure-based drug design.

The experimental evidence is the electron density of the protein and the ligand, and therefore the precision by which we can interpret the electron density and position the atoms determines the structure’s accuracy and reliability. Although there are stringent and strict validation tools for the protein-ligand structure, the PDB has structures with overenthusiastic interpretation of the electron density of the ligand. This can lead to the placement of ligands with no supporting electron density and ultimately affect the accuracy of the structure. Another issue that can arise is a misinterpretation of the ligand identity. This can happen when the electron density map is not clear enough to allow for unambiguous interpretation. Finally, there may be cases where the electron density cannot be fully accounted for, leading to uncertainties in the structure. In conclusion, crystallographic studies of protein-ligand complexes are an important tool in drug discovery, but they require careful interpretation and validation of the obtained results. Here we attempt to discuss some of the pitfalls and tips on improving the accuracy and reliability of the obtained protein-drug structure and ultimately develop better drugs for the treatment of various diseases.

This study is based on our long-term interest in developing new therapies for treating organophosphate (OP) poisoning from pesticides and nerve agents. The enzyme phosphotriesterase (PTE) capable of hydrolyzing OP compounds, including chemical warfare nerve agents and pesticides, has been characterized from a number of bacterial sources, including *Brevundimonas diminuta (Bd)* (formerly *Pseudomonas diminuta*), *Flavobacterium spp*., and *Agrobacterium radiobacter* (Bigley *et al*., 2013; Cherny *et al*., 2013; Goldsmith *et al*., 2016; Harper *et al*., 1988; Horne *et al*., 2002; Masson & Rochu, 2009). The present study focuses on the enzyme from *Bd*_PTE, (Holm & Sander, 1997), a dimeric metallohydrolase that folds as a (β/α)_8_ TIM barrel with two α- and β-Zn^2+^ ions embedded at the active site (Figure 1). Carbon dioxide reacts with the side chain of K169 to form a carbamate functional group within the active site, which serves as a bridging ligand to the two Zn^2+^ ions, both required for full catalytic activity. The α-Zn^2+^ ion is ligated to the protein via direct interactions with H55, H57, and D301, and the β-Zn^2+^ ion with H201, H230, and the carbamate functional group of K169 (Figure 1b).

**Figure 1.**
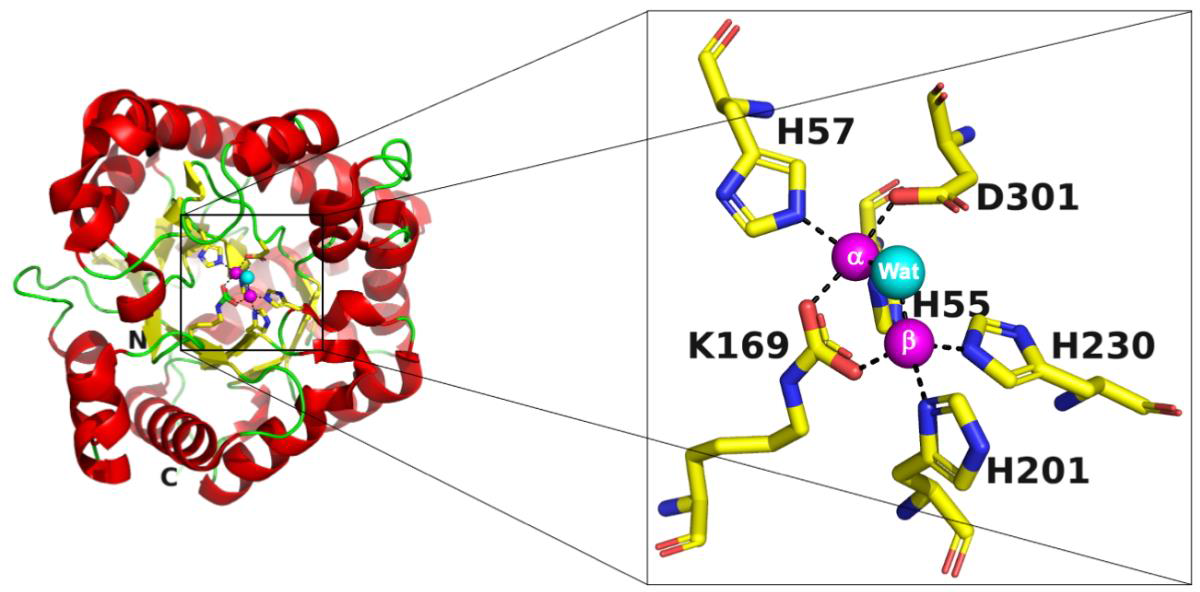
View of a typical PTE structure (PDB-ID 1HZY (Benning *et al*., 2001) (1HZY). (a) the (β/α)_8_ TIM barrel fold is shown as a cartoon, with the helices in red, sheets in yellow, coils in green, and the two α- and β-Zn^2+^ ions as magenta spheres, with a single bridging water shown as a cyan-colored sphere. The six residues that bind to the two Zn^2+^ ions are shown as stick figures with carbon atoms colored yellow, nitrogen atoms colored blue, and oxygen atoms colored red. The N-term and C-term residues are labeled N and C, respectively. (b) close-up view of the active site of the apo-PTE structure (PDB-ID 1HZY). The α-Zn^2+^ ion is directly bound to H55, H57, D301, and the carbamate functional group of K169, while the β-Zn^2+^ ion is bound to H201, H230, and the carbamate functional group of K169. A water molecule forms a bridge between the two Zn ions. The coloring scheme is the same as in Figure 1a.

There has been significant interest in applying PTE (Paraoxonase-like organophosphate hydrolases) for the degradation and disposal of OP-based nerve agents and pesticides (Singh, 2009). Computational design, directed evolution, and site-directed mutagenesis of PTEs have been used extensively over the course of the last decade to enhance the catalytic efficiency, substrate specificity, and stability of PTEs towards a broad range of OPs of interest (Bigley *et al*., 2015; Cherny *et al*., 2013; Goldsmith *et al*., 2017; Goldsmith *et al*., 2016; Grimsley *et al*., 2005; Jackson *et al*., 2009; McLoughlin *et al*., 2005; Tsai *et al*., 2012; Yang *et al*., 2014; Yang *et al*., 2003). These studies have successfully increased the efficiency of PTEs to degrade and detoxify OPs and have shown promising results in enhancing the efficacy of nerve-agent bioscavengers that could serve as prophylactic antidotes for the treatment of both V- and G-type nerve agent intoxication (Cherny *et al*., 2013; Goldsmith *et al*., 2017; Goldsmith *et al*., 2016).

Since the evolved PTE scaffolds had been shown to act as highly potent bioscavengers both *in vitro* and *in vivo*, it was important to clarify the structural features of their active sites that contribute to their efficacy. Until now, there are over fifty OP/PTE crystal structures complexes which have provided valuable information for mapping its active site. We thought it important to determine the crystal structures of our most effective PTE variants and, where possible, complex them with OPs to understand the structural features responsible for their enhanced effectiveness. The following factors were investigated:

- Three molecular PTE variants were chosen based on their high bioscavenging activity: A53, C23 (Cherny *et al*., 2013), and C23M (which corresponds to G5-B60 in Supplement Table 3 (Cherny *et al*., 2013), as well as retaining a residual tag used for protein expression.
- Three different crystallization conditions on A53, C23, and C23M to optimize the formation of PTE/OP complexes.
- Four different space groups to investigate the impact of crystal packing on the formation of PTE/OP complexes.
- Use of four different OP ligands to enable the identification of ligand-specific interactions and differences between PTE variants.

## 2. Materials and methods

### 2.1. Expression and purification of PTE variants

PTE variants were expressed either without any fusion protein or tag, or as a fusion protein with maltose-binding protein (MBP). In the latter case, the MBP was fused to PTE via a Factor Xa cleavage motif (IEGR) and a spacer sequence of eight amino acid residues (ISEFITNS) (see schematic representations in Figure 2a). The purpose of using the MBP fusion partner was to increase PTE’s expression levels and solubility. However, prior to crystallization trials, the MBP fusion partner was removed by digesting it with Factor Xa.

**Figure 2.**
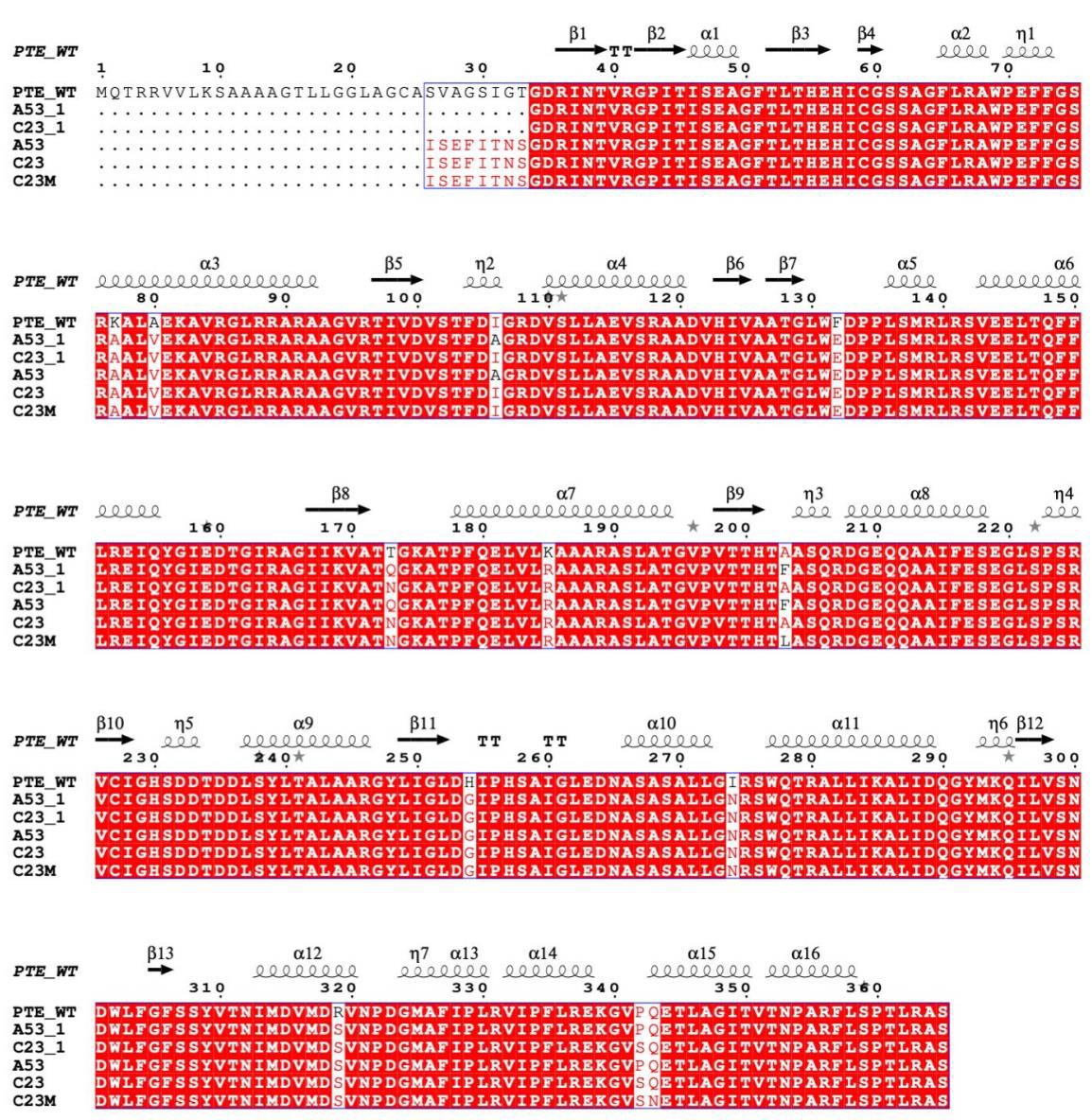
PTE variants (a) Schematic presentation of the maltose-binding protein (MBP) fused before Factor Xa cleavage motif (IEGR) and spacer sequence of eight amino acid residues (ISEFITNS), followed by the mature PTE protein sequence starting with Gly34. Factor Xa cleaves after the Arg residue leaves the ISEFITNS linker, followed by the PTE construct ^34^G---S^365^. (b) Sequence alignment of PTE_WT (PDB-ID 1HZY) (Benning *et al*., 2001), A53_1, C23_1, A53, C23, and C23M. PTE_WT secondary structure elements are labeled above, and that of C23M on the bottom. α-helices and 3_10_-helices (shown with the symbol “η”) are indicated with coils and β-strands with arrows. The residues conserved in all four proteins are in red blocks. The eight amino acid residues linker ^26^IAEFITNS^33^ of the MBP-fused constructs A53 and C23 are shown. Multiple sequence alignment was performed using MultAlin (Corpet, 1988), and the figure was created using ESPript (Robert & Gouet, 2014).

#### 2.1.1. Expression of tag-free PTE variants

PTE constructs, A53, C23, and C23M (Cherny *et al*., 2013) were generated based on a modified protocol (Tokuriki *et al*., 2012) (Figure 2). These variants (residues 34-365), devoid of any tag, were cloned into pET21. They were grown in a 5L culture of *E. coli* BL21 cells at 30°C, following induction with IPTG (0.5mM). The bacterial pellet was resuspended in 20 mM HEPES, pH 7.5, supplemented with 0.1 mM ZnCl_2_. The cells were sonicated for 2 min with 30 s on and 30 s off pulses at 35% amplitude of a Vibracell apparatus (Sonics & Materials, Inc, Newtown, CT). The clarified lysate was loaded onto a HiTrap DEAE_FF 5 ml column (GE Healthcare) pre-equilibrated with 0.1 mM ZnCl_2_/20 mM HEPES, pH 7.5. PTE-containing fractions were eluted with a 0-1 M NaCl gradient in the same buffer. The peak fractions PTE were dialyzed against 0.1 mM ZnCl_2_/50 mM MES, pH 6.0, and loaded onto a Tricorn MonoS 10/100 GL cation exchange column (GE Healthcare) equilibrated with the same buffer. PTE was again eluted with a 0-1 M NaCl gradient in the same buffer. Fractions containing PTE were then concentrated to 5ml and loaded onto a Gel Filtration column (HiLoad Superdex 75 16/60) equilibrated with 50 mM NaCl/25 mM HEPES, pH 8.0. The pooled eluent fractions were applied to a Tricorn Q 10/100 GL anion exchange column equilibrated with 50 mM Tris, pH 8.0, and eluted with a 0-1 M NaCl gradient in the same buffer. The final protein solution was concentrated to 13-17 mg/ml for crystallization screens.

#### 2.1.2. Expression and purification of MBP-PTE fusion proteins

To increase PTE expression levels and solubility, MBP was introduced as a fusion partner to the three PTE variants, with a Factor Xa cleavage motif (IEGR) (Zhao *et al*., 2013) followed by a spacer of 8 amino acids (ISEFITNS) between the two proteins (Figure 2a), and purified as described earlier (Cherny *et al*., 2013). Briefly, BL21 cells were transformed with a PTE plasmid bearing the desired mutation. The BL21 cells were grown overnight at 37°C in LB supplemented with ampicillin. They were then sub-cultured (1% v/v of inoculum) in LB supplemented with 0.2 mM ZnCl_2_ and ampicillin (100µg/ ml) and allowed to grow at 37°C until OD_600 nm_ reached 0.6-0.8, followed by induction with 0.4 mM IPTG and 14-18 h further growth at 20°C. The cell pellet was suspended in buffer A (0.1 mM ZnCl_2_/10 mM NaHCO_3_/100 mM NaCl/100 mM Tris, pH 8.0) containing 0.4 mg/ ml lysozyme, 50 units benzonase, and 1mM PMSF and sonicated using a Vibracell apparatus (Sonics & Materials, Inc, Newtown, CT). The sonication conditions were: 30 sec on, 30 sec off, 30 % amplitude, for 2 min. A clear cell lysate was obtained by centrifugation at 7,500 rpm for 30 min. Amylose beads equilibrated with buffer A were packed into a column. The clarified lysate was loaded onto the column by gravitation and washed with buffer A. The MBP-PTE variants were eluted with 10 mM maltose in buffer A. The eluted fractions were analyzed by 12% SDS-PAGE. Fractions containing MBP-PTE were pooled and dialyzed overnight against buffer A. The protein concentration was measured at 280 nm using a molar extinction coefficient for MBP-PTE of 95,800 cm^-1^ M^-1^.

For crystallization trials, A53, C23, and C23M variants were expressed and purified from 2.5-5L cultures as above. The suspended pellet was disrupted using a cell disruptor (Constant Systems Ltd, Low March, UK). Following elution of the fusion protein from the amylose beads followed by dialysis to remove maltose, MBP was cleaved from the PTE variant by incubation with 25 μg per 800 ml culture of factor Xa (Zhao *et al*., 2013) for 40 h at 4°C. Post cleavage, the solution was passed 4 times over a column packed with fresh amylose beads to retain the cleaved MBP and any residual uncleaved MBP-PTE, while tagless PTE appeared in the flowthrough. In some cases in which PTE was still contaminated with residual MBP, an anion exchange column was also used to separate the tagless PTE from residual MBP. In these cases, the protein solution was dialyzed against 0.1mM ZnCl_2_/10% glycerol/200mM NaCl/20mM Tris, pH 8.0, and loaded onto a Tricorn Q10/100 column (GE Healthcare). Under these conditions, pure PTE came out in the flowthrough of the column, while MBP was retained. The tagless PTE was dialyzed against buffer A and concentrated to ∼11 mg/ml.

### 2.2. Crystallization and structure determination of PTE variants

Crystals of the three PTE variants, *i.e.*, A53, C23, and C23M, were grown at 19°C by the hanging drop vapor diffusion technique using a Mosquito robot (TTP LabTech, Inc., Cambridge, MA). The crystals of the PTE variants obtained in the presence of three different crystallization precipitates, including (1) Polyethylene glycol (PEG) 6000 and 2-methyl-2,4-pentanediol (MPD), (2) glycerol and (NH_4_)_2_SO_4_ (AS) and (3) Polyacrylic acid (PAA). The crystals grew in four different space groups P2_1_2_1_2, P2_1_2_1_2_1_, P2_1_, and P4_3_2_1_2. All crystals (but one) diffracted to high resolution in the 1.38-2.0 Å range. Crystallization optimization experiments were set up manually by the hanging drop vapor diffusion technique. Four phosphonate ligands were co-crystallized with A53, C23, and C23M variants (Figure 3). Twelve crystal structures were analysed, including the apo A53, C23, and C23M variants, as well as several OP ligands bound to the PTE variants.

**Figure 3.**
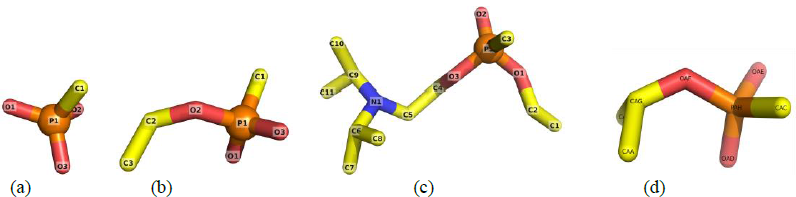
OP ligands co-crystallized with PTE variants. (a) Methylphosphonate (I); (b) O-ethyl methylphosphonate (II); (c) O-ethyl-O-(N,N-diisopropylaminoethyl) methylphosphonate (III) (oxo analog of VM); (d) O-isopropyl methylphosphonate.

X-ray data collections were performed under cryo conditions (100 K), either in-house on a Rigaku RU-H3R X-ray or on beamlines ID14-4 and ID23-1 at the ESRF (Grenoble, France) (Table 1). All diffraction images were indexed and integrated using the program Mosflm, (Leslie & Powell, 2007), and the integrated reflections were scaled using the SCALA program (Evans, 2006). Structure factor amplitudes were calculated using TRUNCATE (French & Wilson, 1978) from the CCP4 program suite. The structure of the apo-A53 (A53_1) was solved by molecular replacement with the program PHASER (McCoy *et al*., 2007) using the homologous refined structure of the Bd_PTE as a model (PDB-ID 1HZY (Benning *et al*., 2001) (1HZY). All steps of atomic refinements were carried out with the CCP4/REFMAC5 program (Murshudov *et al*., 1999) and Phenix (Afonine *et al*., 2012). The models were built into the electron density map using the COOT program (Emsley & Cowtan, 2004). Following this, the refined A53_1 structure was used as a model to determine the structure of all the other PTE variants (Table 1). All models were optimized using the PDB-REDO (Joosten *et al*., 2014) and were evaluated with MolProbity (Chen *et al*., 2010). Details of the refinement statistics of the PTE’s structures are described in Table 1.

**Table 1.** Crystallization, data collection and refinement statistics for the PTE’s

| Crystal | A53_1 | A53_2 | A53_3 |
| --- | --- | --- | --- |
| Ligand in the active site | Apo<br>One Zn | Apo<br>Q5K | Apo<br>D9K |
| **Crystallization conditions** | 12% PEG 6,000<br>5% MPD<br>0.1M HEPES pH=7.5 | 1.5M (NH<sub>4</sub>)<sub>2</sub>SO<sub>4</sub><br>17% Glycerol<br>0.1M Tris pH=7.5 | 16% PEG 6,000<br>2% MPD<br>0.1M HEPES pH=7.5<br>0.1M Glycine |
| **Data Collection** |  |  |  |
| Data collected at | ID14-4 ESRF | Rigaku home source | Rigaku home source |
| PDB code | 8P7F | 8P7H | 8P7I |
| Space group | *P2<sub>1</sub>2<sub>1</sub>2* | *P4<sub>3</sub>2<sub>1</sub>2* | *P2<sub>1</sub>* |
| Cell dimensions: |  |  |  |
| a,b,c (Å) | 79.70 93.67 44.59 | 69.67 69.67 186.83 | 54.23 81.25 70.66 |
| α,β,γ (°) | 90, 90, 90 | 90, 90, 90 | 90, 94.80, 90 |
| No. of copies in a.u. | 1 | 1 | 2 |
| Resolution (Å) | 39.85-2.00 | 25.48-1.77 | 36.77-1.70 |
| Upper resolution shell (Å) | 2.071-2.00 | 1.84-1.77 | 1.76-1.70 |
| Unique reflections | 23,037 (2,273) | 44,626 (4,284) | 67,098 (6,633) |
| Completeness (%) | 98.91 (99.91) | 97.93 (95.84) | 99.69 (98.94) |
| Average I/σ(I) | 12.50 (4.2) | 22.9 (7.4) | 20.06 (4.06) |
| Multiplicity | 5.3 (5.5) | 17.0 (16.7) | 6.1 (5.8) |
| CC1/2 | 0.727 (0.638) | 0.952 (0.91) | 0.925 (0.925) |
| R-pim | 0.218 (0.306) | 0.086 (0.141) | 0.024 (0.176) |
| **Refinement** |  |  |  |
| Resolution range (Å) | 39.85-2.00 | 25.48-1.77 | 36.77-1.70 |
| No. of reflections (I/σ(I) > 0) | 23,010 (2,271) | 44,611 | 67,085 |
| No. of reflections in test set | 2,000 (197) | 4,283 | 3,397 |
| R-working / R-free | 0.2014 / 0.2297 | 0.1574 / 0.1894 | 0.1511 / 0.1808 |
| No. of protein atoms | 2163 | 2535 | 5067 |
| No. of water molecules | 86 | 401 | 537 |
| Overall average B factor (Å<sup>2</sup>) | 41.93 | 20.23 | 25.16 |
| Root mean square deviations: |  |  |  |
| - bond length (Å) | 0.007 | 0.006 | 0.006 |
| - bond angle (°) | 0.85 | 0.85 | 0.85 |
| **Ramachandran Plot** |  |  |  |
| Most favored (%) | 97.50 | 96.94 | 97.58 |
| Additionally allowed (%) | 2.14 | 3.06 | 2.42 |
| Disallowed (%) | 0.36 | 0.00 | 0.00 |

\* Values in parentheses refer to the data of the corresponding upper resolution shell

| **Crystal** | A53_4 | A53_5 |
| --- | --- | --- |
| Ligand in the active site | Apo<br>Polyacrylic acid | Methylphosphonate |
| **Crystallization conditions** | 22% Polyacrylic acid<br>0.02M MgCl<sub>2</sub><br>0.1M HEPES pH=7.5 | 1.5M (NH<sub>4</sub>)<sub>2</sub>SO<sub>4</sub><br>12% Glycerol<br>0.1M Tris pH=7.5 |
| **Data Collection** |  |  |
| Data collected at | Rigaku home source | Rigaku home source |
| PDB code | 8P7K | 8P7M |
| Space group | *P2<sub>1</sub>* | *P4<sub>3</sub>2<sub>1</sub>2* |
| Cell dimensions: |  |  |
| a,b,c (Å) | 55.03 81.49 70.81 | 69.74 69.74 186.76 |
| α,β,γ (°) | 90, 94.99, 90 | 90, 90, 90 |
| No. of copies in a.u. | 2 | 1 |
| Resolution (Å) | 32.37-1.93 | 27.94-1.85 |
| Upper resolution shell (Å) | 2.00-1.93 | 1.92-1.85 |
| Unique reflections | 46,379 (4,410) | 39,668 (3,811) |
| Completeness (%) | 98.69 (93.67) | 98.24 (96.43) |
| Average I/σ(I) | 10.80 (2.16) | 36.19 (11.23) |
| Multiplicity | 3.5 (2.9) | 17.2 (17.0) |
| CC1/2 | 0.995 (0.772) | 1 (0.989) |
| R-pim | 0.052 (0.338) | 0.014 (0.063) |
| **Refinement** |  |  |
| Resolution range (Å) | 32.37-1.93 | 27.94-1.85 |
| No. of reflections (I/σ(I) > 0) | 46,363 | 39,642 |
| No. of reflections in test set | 2,355 | 3,809 |
| R-working / R-free | 0.1593 / 0.2119 | 0.1612 / 0.1876 |
| No. of protein atoms | 5015 | 2506 |
| No. of water molecules | 463 | 313 |
| Overall average B factor (Å<sup>2</sup>) | 27.32 | 18.52 |
| Root mean square deviations: |  |  |
| - bond length (Å) | 0.007 | 0.006 |
| - bond angle (°) | 0.83 | 0.92 |
| **Ramachandran Plot** |  |  |
| Most favored (%) | 97.25 | 97.24 |
| Additionally allowed (%) | 2.75 | 2.76 |
| Disallowed (%) | 0.0 | 0.00 |

| Crystal | C23_1 | C23_2 | C23_3 |
| --- | --- | --- | --- |
| Ligand in the active site | Apo | O-isopropyl methylphosphonate | Methylphosphonate |
| **Crystallization conditions** | 10% PEG 6,000<br>2% PEG 400<br>15% MPD<br>0.1M HEPES pH=7.5 | 1.5M (NH<sub>4</sub>)<sub>2</sub>SO<sub>4</sub><br>17% Glycerol<br>0.1M Tris pH=7.0 | 1.5M (NH<sub>4</sub>)<sub>2</sub>SO<sub>4</sub><br>17% Glycerol<br>0.1M Tris pH=8.5 |
| **Data Collection** |  |  |  |
| Data collected at | ID14-4 ESRF | Rigaku home source | Rigaku home source |
| PDB code | 8P7N | 8P7Q | 8P7R |
| Space group | *P2<sub>1</sub>2<sub>1</sub>2<sub>1</sub>* | *P4<sub>3</sub>2<sub>1</sub>2* | *P4<sub>3</sub>2<sub>1</sub>2* |
| Cell dimensions: |  |  |  |
| a,b,c (Å) | 88.89 119.90 161.49 | 69.56 69.56 186.64 | 69.83 69.83 186.78 |
| α,β,γ (°) | 90, 90, 90 | 90.0, 90.0, 90.0 | 90, 90, 90 |
| No. of copies in a.u. | 4 | 1 | 1 |
| Resolution (Å) | 77.78-3.20 | 34.78-1.77 | 28.43-1.85 |
| Upper resolution shell (Å) | 3.31-3.20 | 1.87-1.77 | 1.92-1.85 |
| Unique reflections | 29,131 (2,870) | 45,349 (4,453) | 40,443 (3,956) |
| Completeness (%) | 99.61 (99.90) | 99.92 (100) | 99.94 (100.00) |
| Average I/σ(I) | 7.24 (2.64) | 18.97 (6.41) | 33.96 (5.21) |
| Multiplicity | 4.1 (4.2) | 20.4 (20.6) | 12.0 (11.7) |
| CC1/2 | 0.987 (0.823) | 0.999 (0.974) | 0.859 (0.689) |
| R-pim | 0.09627 (0.3261) | 0.0241 (0.0978) | 0.1528 (0.2906) |
| **Refinement** |  |  |  |
| Resolution range (Å) | 71.41-3.20 | 31.11-1.77 | 28.43-1.85 |
| No. of reflections (I/σ(I) > 0) | 29,063 | 45,340 | 40,437 |
| No. of reflections in test set | 1,412 | 4,453 | 2,015 |
| R-working / R-free | 0.1830 / 0.2546 | 0.167 / 0.191 | 0.1548 / 0.1782 |
| No. of protein atoms | 9890 | 2499 | 2541 |
| No. of water molecules | 8 | 259 | 377 |
| Overall average B factor (Å<sup>2</sup>) | 44.15 | 19.17 | 17.54 |
| Root mean square deviations: |  |  |  |
| - bond length (Å) | 0.010 | 0.016 | 0.006 |
| - bond angle (°) | 1.21 | 2.000 | 0.92 |
| **Ramachandran Plot** |  |  |  |
| Most favored (%) | 95.69 | 96.34 | 97.26 |
| Additionally allowed (%) | 4.08 | 3.35 | 2.74 |
| Disallowed (%) | 0.23 | 0.30 | 0.00 |

| **Crystal** | C23_4 | C23_5 |
| --- | --- | --- |
| Ligand in the active site | O-ethyl-O-(N,N-diisopropylaminoethyl) methylphosphonate<br>Hydrolyzed | O-Ethyl methylphosphonate |
| **Crystallization conditions** | 1.0M (NH<sub>4</sub>)<sub>2</sub>SO<sub>4</sub><br>12% Glycerol<br>0.1M Tris pH=7.5 | 1.5M (NH<sub>4</sub>)<sub>2</sub>SO<sub>4</sub><br>17% Glycerol<br>0.1M Tris pH=8.5 |
| **Data Collection** |  |  |
| Data collected at | Rigaku home source | Rigaku home source |
| PDB code | 8P7S | 8P7T |
| Space group | *P4<sub>3</sub>2<sub>1</sub>2* | *P4<sub>3</sub>2<sub>1</sub>2* |
| Cell dimensions: |  |  |
| a,b,c (Å) | 69.54 69.54 185.84 | 69.45 69.45 186.97 |
| α,β,γ (°) | 90, 90, 90 | 90, 90, 90 |
| No. of copies in a.u. | 1 | 1 |
| Resolution (Å) | 33.77-1.77 | 23.19-1.80 |
| Upper resolution shell (Å) | 1.84-1.77 | 1.86-1.80 |
| Unique reflections | 45,145 (4,428) | 42,215 (4,018) |
| Completeness (%) | 99.89 (99.84) | 97.23 (94.96) |
| Average I/σ(I) | 30.97 (12.07) | 22.59 (5.53) |
| Multiplicity | 11.8 (11.4) | 20.4 (20.6) |
| CC1/2 | 0.888 (0.914) | 0.999 (0.948) |
| R-pim | 0.09951 (0.121) | 0.02038 (0.1358) |
| **Refinement** |  |  |
| Resolution range (Å) | 31.10-1.77 | 22.97-1.80 |
| No. of reflections (I/σ(I) > 0) | 45,111 | 42,212 |
| No. of reflections in test set | 4,428 | 4,107 |
| R-working / R-free | 0.1602 / 0.1841 | 0.1569 / 0.1844 |
| No. of protein atoms | 2540 | 2522 |
| No. of water molecules | 346 | 268 |
| Overall average B factor (Å<sup>2</sup>) | 18.88 | 20.07 |
| Root mean square deviations: |  |  |
| - bond length (Å) | 0.006 | 0.006 |
| - bond angle (°) | 0.92 | 1.03 |
| **Ramachandran Plot** |  |  |
| Most favored (%) | 97.85 | 97.26 |
| Additionally allowed (%) | 2.15 | 2.74 |
| Disallowed (%) | 0.00 | 0.00 |

| **Crystal** | C23M_1 | C23M_2 |
| --- | --- | --- |
| Ligand in the active site | Apo- Zn wavelength<br>Z7K | Methylphosphonate |
| **Crystallization conditions** | 1.5M (NH<sub>4</sub>)<sub>2</sub>SO<sub>4</sub><br>12% Glycerol<br>0.1M Tris pH=8.0 | 1.5M (NH<sub>4</sub>)<sub>2</sub>SO<sub>4</sub><br>12% Glycerol<br>0.1M Tris pH=8.0 |
| **Data Collection** |  |  |
| Data collected at | ID23_1 ESRF | Rigaku home source |
| PDB code | 8P7U | 8P7V |
| Space group | *P*4<sub>3</sub>2<sub>1</sub>2 | *P*4<sub>3</sub>2<sub>1</sub>2 |
| Cell dimensions: |  |  |
| a,b,c (Å) | 69.69 69.69 187.07 | 69.70 69.70 186.48 |
| α,β,γ (°) | 90, 90, 90 | 90, 90, 90 |
| No. of copies in a.u. | 1 | 1 |
| Resolution (Å) | 18.11-1.38 | 25.46-1.74 |
| Upper resolution shell (Å) | 1.42-1.38 | 1.80-1.74 |
| Unique reflections | 96,151 (9,438) | 47,579 (4,555) |
| Completeness (%) | 99.15 (98.70) | 98.25 (96.36) |
| Average I/σ(I) | 6.87 (2.03) | 20.05 (3.92) |
| Multiplicity | 6.5 (6.4) | 20.0 (20.0) |
| CC1/2 | 0.992 (0.776) | 0.999 (0.895) |
| R-pim | 0.05118 (0.274) | 0.02354 (0.2009) |
| **Refinement** |  |  |
| Resolution range (Å) | 18.11-1.38 | 24.88-1.74 |
| No. of reflections (I/σ(I) > 0) | 95,540 | 47,567 |
| No. of reflections in test set | 4,684 | 2,326 |
| R-working / R-free | 0.1572 / 0.1781 | 0.1668 / 0.1886 |
| No. of protein atoms | 2557 | 2506 |
| No. of water molecules | 372 | 305 |
| Overall average B factor (Å<sup>2</sup>) | 19.53 | 20.12 |
| Root mean square deviations: |  |  |
| - bond length (Å) | 0.066 | 0.007 |
| - bond angle (°) | 0.95 | 1.02 |
| **Ramachandran Plot** |  |  |
| Most favored (%) | 96.34 | 97.86 |
| Additionally allowed (%) | 3.66 | 1.83 |
| Disallowed (%) | 0.00 | 0.31 |

## 3. Results

### 3.1. The correlation between the presence of Zn^+2^ and the absence of tags: A53_1 and C23_1

The A53 variant, devoid of any tags, was crystallized in the presence of 12% PEG 6000, 5% MPD, and 0.1M HEPES, pH 7.5, and diffracted to 2.0 Å (A53_1 in Table 1). The (β/α)_8_ TIM barrel fold of A53_1 is very similar to that of WT *Bd*_PTE, PDB-ID 1PTA (1PTA) (Benning *et al*., 2001) (Figure 4). However, the A53_1 is a monomer with only one Zn^2+^ ion in the active site. In contrast, virtually all PTEs require two divalent metals in their active site center as they contribute to the activity and stability of the enzyme (Arnold & Haymore, 1991). Removal of any of the two bound Zn^2+^ ions can lead to a loss of enzymatic activity. The α-Zn^2+^ ion in the PTE known structures is ligated with H55, H57, and D301, and the β-Zn^2+^ ion coordinated to H201, H230, and the carbamate functional group bound to K169 (Figure 1b).

**Figure 4.**
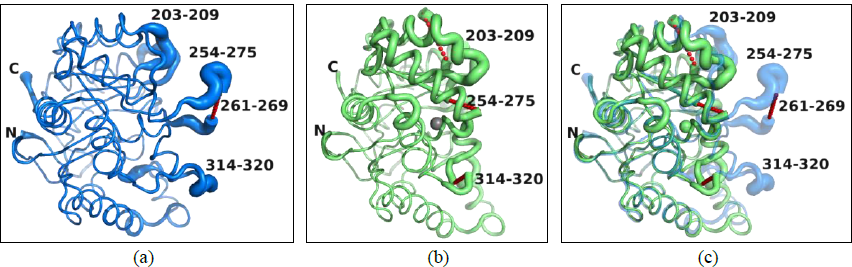
Cartoon putty representation of the (β/α)_8_ TIM barrel fold of *Bd*_PTE structures, with regions of larger temperature factors as fatter lines and those with smaller temperature factors as thinner lines. (a) 1PTA structure (Benning *et al*., 1994) with no Zn ions, with missing residues, 261-269 shown in a dashed red trace; (b) A53_1 with one Zn^+2^ ion, shown in the same orientation as in (a) with the three regions of residues not seen in the electron density, *i.e.*, 203-209, 254-275 and 314-320, labeled; (c) Superposition of the A53_1 on 1PTA, in the same orientation as in (a) showing how similar they are except for the missing residues in A53_1, with A53_1 containing one Zn ion shown in green, and 1PTA shown in magenta. The regions of high-temperature factors fall into similar regions in these two structures. The 1PTA structure is shown as 50% transparent to see the missing regions in A53_1 more easily.

The A53_1 and the 1PTA crystallize in the same P2_1_2_1_2 Space Group with virtually identical unit cells a=79.70Å, b=93.67Å c=44.59Å, and a=80.2Å, b=93.7Å, c=45Å, respectively. It is interesting to note that the 1PTA structure does not contain a Zn ion in its active site (Figure 5a), despite being structurally similar to the A53_1 variant (RMS deviation of a 0.498Å), which contains one Zn ion in its active site (Figure 5b). The putative active sites of 1PTA and A53_1 structures differ significantly from the canonical active site containing two Zn ions, as seen in other PTE structures such as 1HZY (Figure 5c).

**Figure 5.**
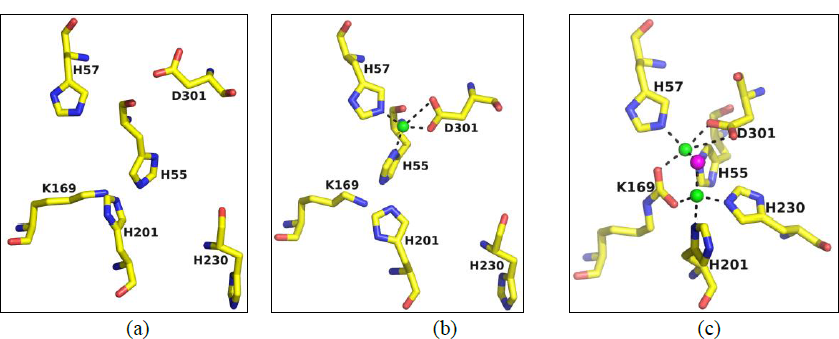
Comparison of the active site regions of PTEs with different numbers of Zn ions bound. (a) WT *Bd*_PTE (PDB-ID 1PTA) with no Zn^2+^ ions; (b) *Bd*_PTE (A53_1) with one Zn^2+^ ion and (c) *Bd*_PTE (PDB-ID 1HZY) with two Zn^2+^ ions.

It has been reported that some reagents used in the purification and crystallization processes can act as chelators, creating coordinate bonds with the Zn^2+^ ions in solution thereby decreasing their effective concentration (Fischer *et al*., 1979). In this study, the Tris buffer was used in high concentrations (>50mM) in the purification and some of the crystallization experiments of the PTE variants. Tris has metal-binding properties, especially to Zn^2+^ ions. The amine nitrogen atom in Tris can chelate Zn^2+^ ions, decreasing the effective concentration of free Zn^2+^, making it difficult to incorporate two Zn^2+^ ions into the binding site of the PTE variants. Since the presence of two Zn^2+^ ions and the correct orientation of the residues in the active site are crucial for maintaining PTE’s catalytic activity, it is reasonable to assume that the A53_1 monomer is catalytically inactive. To avoid forming non-active PTE molecules containing less than two Zn^2+^ ions in the active site, it was crucial to add ZnCl_2_ during protein expression, purification, and crystallization. All the structures in Table 1, except A53_1 and C23_1, were prepared this way and exhibited a dimer with two Zn^2+^ ions in the active site.

In comparing the *Bd*_PTE WT structures, 1HZY which contains two Zn^2+^ ions, with A53_1 (monomer), they have an RMS deviation of 0.65 Å. However, inspection of the active site of A53_1 showed that the α-Zn^2+^ ion is ∼2.5 Å away from the α-Zn^2+^ ion in the 1HZY structure, while the β-Zn^2+^ is not observed at all in the A53_1 structure (Figure 5). It appears, in fact, that A53_1 more closely resembles the structure of 1PTA white no Zn^2+^ ions (Figure 5). There is also a sizeable conformational deviation of key active site residues, with the NH_2_ group of the K169 side chain pointing in the opposite direction (2.2 Å away), and the carbamate moiety is absent compared to the 1HZY structure. Thus, in the A53_1 structure, K169 is not interacting with the two Zn^2+^ ions. The imidazole ring of H230 is flipped by 180°, pointing away from the putative β-Zn^2+^ ion, and the imidazole ring of H201 adopts a different orientation as compared to that in the 1HZY structure. These conformational changes are most likely due to the absence of the β-Zn^2+^ ion, to which the side chains of H201 and H230 are expected to coordinate.

While virtually all PTEs in the PDB are seen to crystallize as molecular dimers in the asymmetric unit, these two structures, A53_1 and 1PTA, show only one molecule in the asymmetric unit. They each form a dimer by applying the crystallographic 2-fold axis in the P2_1_2_1_2 space group. However, this dimer interface utilizes different residues and is rather loose compared to the canonical non-crystallographic dimer seen in PTEs, suggesting that it is not a physiological dimer (Figure 8). The two segments, residues 60-79 and 301-313, involved in the canonical PTE dimer interface show pronounced conformational differences in the A53_1 structure cartoon in (Figure 4a). Furthermore, three regions, 203-209, 254-275, and 314-320, in the A53_1 structure are disordered and thus not visible in the electron density map in Figure 4a. The latter (314-320) is in the vicinity of residues 301-313 that are involved in the canonical PTE dimer interface. Since some of the residues in the disordered regions are in the vicinity of the PTE active site, it is plausible that their disorder and the significant conformational changes observed in residues 60-79 and 301-313 eliminate the activity and precludes the dimerization of A53_1 structure. Indeed, the A53_1 sample is a monomer also in solution, as the Gel Filtration indicates. As mentioned above, to avoid the formation of non-active PTE monomers, ZnCl_2_ was added during the process of protein expression, purification, and crystallization in all the structures except A53_1 and C23_1 (see Table 1).

The C23 tag-free variant was expressed in its native form, without the MBP fusion tag, and purified by conventional purification techniques. The crystals of C23_1 obtained from 10% PEG 6000, 15% MPD, 2% PEG 400, and 0.1M HEPES, pH 7.5, diffracted to 3.2 Å (Table 1). No unidentified ligand was observed in the active site, as seen in A53_2, A53_3, and C23M_1 (see below), however, the C23_1 crystal diffracted very poorly and yielded poor crystal structure and therefore made it unsuitable for use in the OP co-crystallization experiments.

It is interesting to note that the C23_1 crystal structure (tag-free construct. Figure 2) displayed the canonical PTE dimer with two Zn^+2^ ions in each monomer, despite being crystallized under similar conditions as A53_1 without any Zn^+2^ ions added in any of the steps (Figure 6). This suggests that the C23 variant may have a higher affinity for Zn^+2^ than the A53 variant, possibly due to small differences in their amino acid sequences. It is plausible that C23 construct could bind traces of Zn^+2^ more efficiently and retain them more effectively during purification, which prevented the formation of a non-active monomer.

**Figure 6.**
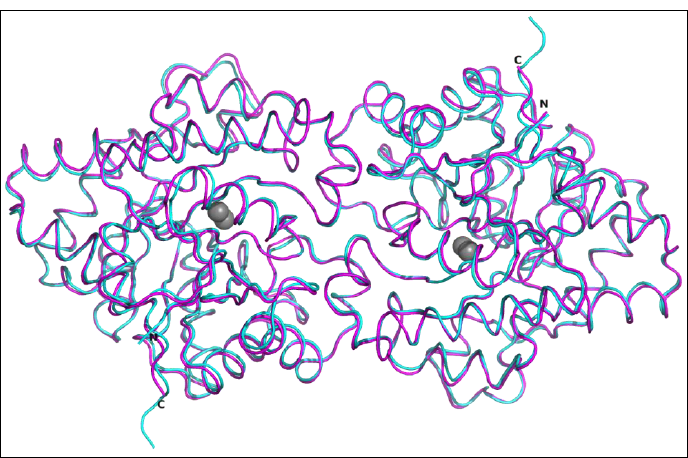
Comparison of the dimer structure of C23_1 (magenta) versus 1HZY (cyan). The two active site Zn^2+^ ions are shown as grey spheres. The two structures are virtually identical.

### 3.2. The structures of tagged constructs of A53, C23, and C23M

The addition of ZnCl_2_ during expression, purification, and crystallization resulted in the presence of two Zn^2+^ ions in the active site of A53_2, A53_3, and A53_4 structures, all crystallized in different crystallization conditions. The coordination geometry of the two Zn^2+^ ions in the active site of these structures is similar to that observed in other PTE structures, such as 1HZY. Interestingly, the crystal structure of A53_2 has a monomer in the asymmetric unit (Figure 7), and the dimer is formed through a crystallographic 2-fold axis in the P4_3_2_1_2 space group as observed in the structure of other PTE variants (A53_5, C23_2, C23_3, C23_4, C23_5, C23M_1, and C23M_2). While the A53_3 and A53_4 crystalize in the P2_1_ space group with a dimer in the asymmetric unit (Figure 8 and Table 1).

**Figure 7.**
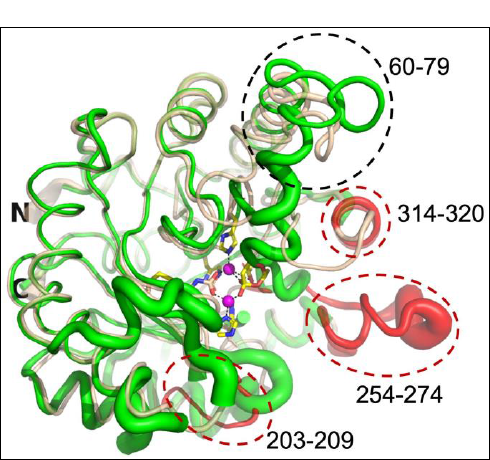
Cartoon tube diagrams of the backbones of the monomer structures of Apo A53_2 (beige) and Apo A53_1 (green). The sizeable conformational difference (residues 60-79) is enclosed in a black dashed line. This region participates in the dimer formation of A53_2, A53_3, and A53_4 structures, with the conformation of A53_2 being very similar to A53_3 and A53_4 but not to that of A53_1. The other missing regions in A53_1, *i.e.*, 203-209, 254-274, and 314-320, are colored red and enclosed in a red dash line. The two Zn^+2^ ions are shown as magenta spheres, and the residues binding to them are shown as sticks.

**Figure 8.**
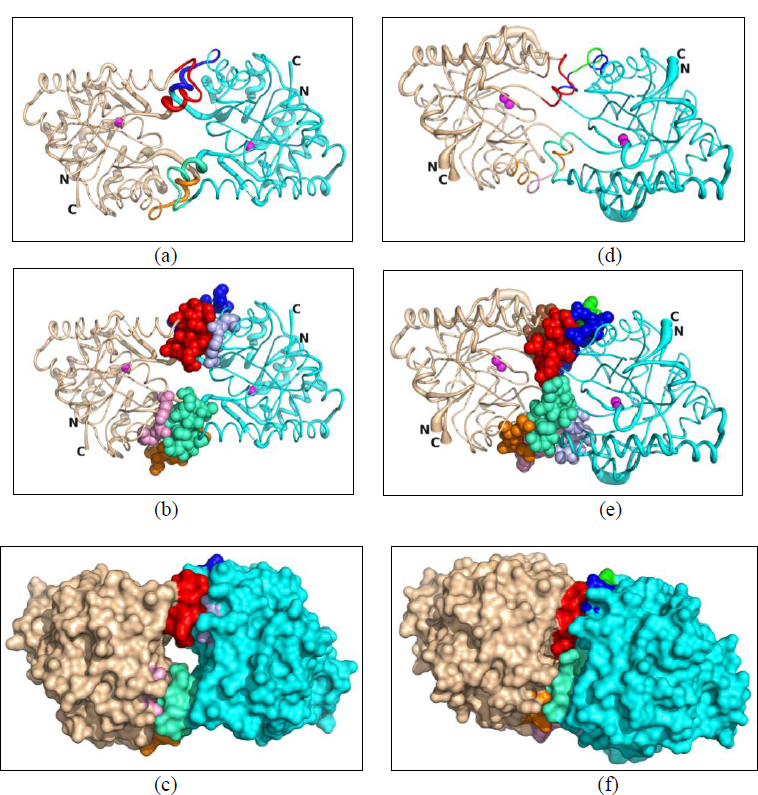
Comparison of the dimer interfaces in Apo A53_1 (left) and Apo A53_2 (right), with subunits A and B shown in beige and cyan. Apo A53_1 is shown with its 2-fold crystallographic symmetry mate in the P4_3_2_1_2 space group. The residues involved in forming the rather loose dimer of A53_1 consist of residues 62-73 (in red) of subunit A and residues 137-144 (blue) of subunit B, as well as residues 152 and 159 of subunit A and residues 67-71 of subunit B, as well as the symmetry-related residues of B to A. For A53_2, the contacts are much tighter, forming the dimer consisting of three regions: residues 62-71 (in red) of monomer A with 137-148 & 152 (blue) of monomer B; residues 133-136 of monomer A with 133-139 of monomer B (those interactions overlap with the 136-152 interactions, so they are not highlighted); and residues 141-145 (in pink) of monomer A with residues 307-311 (in light blue) from monomer B, as well as the symmetry-related residues of B to A. It is interesting to note that region 133-139 of one monomer interacts with 3 regions of the other monomer (62-71, 133-148, and 307-311). (a) A53_1 cartoon trace shown in putty format (thickness of the trace is a function of the B-factors); (b) as in (a) with atoms in residues involved in inter-subunit packing shown as spheres; (c) surface representation of the packing subunits; (d) as in (a) for A53_2; (e) as in (b) for A53_2; (f) as in (c) for A53_2.

### 3.3 Impact of a residual tag on Apo structures of A53, C23, and C23M variants

To facilitate purification, the A53, C23, and C23M variants were expressed as constructs with an N-terminal MBP tag fused before a Factor Xa cleavage motif (IEGR) and an 8-amino acid linker (ISEFITNS), followed by the mature PTE protein sequence starting with Gly34 (MBP-Factor Xa-8 amino acid linker-PTE) (Figure 2a). This resulted in proteins with an 8-residue tag at the N-terminus (Figure 2b).

Apo structures of the 8-residue tagged A53 variant were obtained using three different crystallization precipitants: AS with glycerol, PEG 6000 with MPD, and PAA, which resulted in A53_2, A53_3, and A53_4, respectively. Crystals of the apo C23 were obtained using two precipitants: PEG 6000 with MPD (C23_1), PAA, and AS with glycerol (data not shown).

The C23_1 crystallized in space group P2_1_2_1_2_1_ with two dimers in the asymmetric unit. The crystals of tagged C23 (C23_2, C23_3, C23_4, and C23_5) and C23M (C23M_1 and C23M_2) grew from AS and Glycerol, with one monomer in the P4_3_2_1_2 space group such that the canonical dimer is formed by the crystallographic symmetry.

The unexpected observation was made in the crystal structures of A53_3 and A53_4, where electron density corresponding to residues from the octapeptide spacer ^26^ISEFITNS^33^ was observed. This peptide was left following cleavage of the Factor Xa motif, removing the MBP. The residual peptide of subunit B was found to penetrate the active site of the symmetry-related subunit A, such that the distance between the active site of the A subunit and the N-terminus (I26) of the octapeptide of subunit B was approximately 4.5 Å in A53_3 (Figure 9). The residues from the octapeptide of subunit B made contact with W131, E132, Q173, F203, A270, F306, S308, and Y309 amino acid residues of the symmetry-related subunit A. Interestingly, although both A53_3 and A53_4 crystals were obtained from different crystallization precipitates (PEG 600 with MPD and PAA respectively) they both crystallized in the P2_1_ space group and both had the residual peptide. Since the residual octapeptide was observed only in the P2_1_ space group, it is reasonable to assume that the penetration of the residual octapeptide to the active site is space-group dependent. Moreover, the presence of the octapeptide in the active site could interfere with the binding of the OPs and thus likely explains why no OPs were found in the crystals of A53_3 and A53_4 crystallized in the P2_1_ space group (see Figure 9).

**Figure 9.**
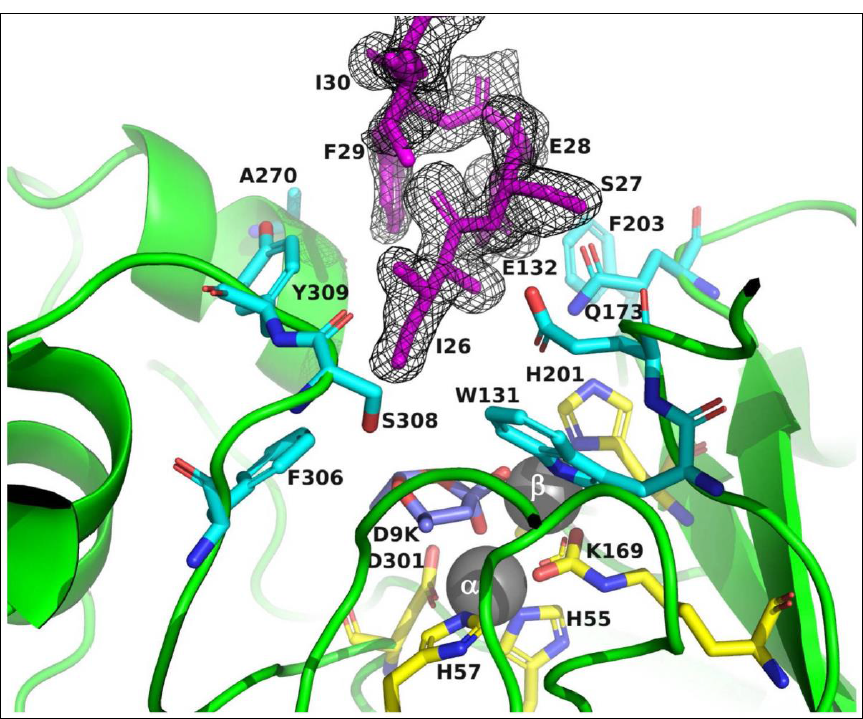
A53_3 shows the 8-residue tag on subunit B entering into the active site region of a symmetry-related subunit A. The tag is shown in magenta. The active site residues of the symmetrically related Chain A are shown in yellow, while those residues within 5 Å of the tag are shown in cyan. Given that the N terminal tag residues are in close contact with active site residues of a symmetry-related molecule, they could behave as a mimic peptide inhibitor and thus may help define the mode of binding of substrates viz. V-type nerve agents.

### 3.4. PTE crystal structures obtained from Polyacrylic acid (PAA)

The PTE variants were also crystallized from Polyacrylic acid (PAA) compound, A53 (A53_4 Table 1), C23, and C23M (data not shown). In these cases, no water molecules are directly bound to Zn^2+^ ions as the acrylic acid used in the crystallization solution was detected in the active site. The bound acrylic acid act as a ligand to bridge the two Zn^2+^ metal ions, mimicking the tetrahedral intermediate formed during the hydrolysis of carboxylate esters (Figure 10). The presence of crystallization reagents in the active site is not unusual, as other PTE structures have also been observed with bound compounds, *e.g.*, cacodylate buffer (PDB-ID 4XD4 and 3RHG) with its two oxygen atoms binding the two Zn^2+^ ions. In the structure of organophosphorus acid anhydrolase (OPAA) (PDB-ID 4ZWO) the glycolic acid similar to the acrylic acid was observed in the active site. Much work has been published on nasal drug delivery with polyacrylic derivatives, and the bioadhesive properties of PAA are well-recognized. PAA is known to have strong bioadhesive properties and a high capacity to bind proteins (Dai *et al*., 2006) which could explain its presence in the active site of the PTE variants. Notably, no electron density for the co-crystallized OP ligands was observed in any PTE variants crystallized from PAA. This suggests that the PAA may have a higher binding affinity to the Zn^+2^ ions compared to OP ligands. Furthermore, the presence of the PAA in the active site in the crystal structure of A53_4 (described in section 2.3.3) suggests that the crystallization reagent, PAA, initially binds to the PTE active site in solution and subsequently, the octapeptide spacer ^26^ISEFITNS^33^ penetrates the active site upon crystal formation in the P2_1_ space group.

**Figure 10.**
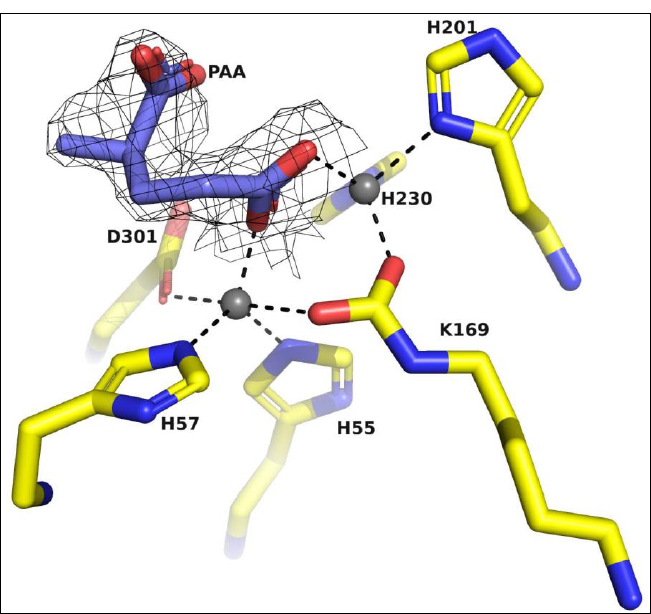
A53_4 showing PAA bound in the active site preventing any OPs from binding.

### 3.5. Identifying the metal type in the PTE metallohydrolase active site

Identifying the intrinsically bound metal ions in a protein-metal complex structure is crucial to ensure that they are consistent with the solutions used during the protein expression, purification, and crystallization steps (Zheng et al., 2008). Unexpected metals could potentially replace the expected metal ions and result in functional differences. In the case of the PTE variants, it was essential to confirm the identity of metal ions in the active site. Therefore, the C23M_1 X-ray data were collected at the Zn absorption edge wavelength (λ=1.2724 Å on beamline ID23_1 at ESRF) (Figure 11), on a crystal that diffracted to 1.38 Å (C23M_1 in Table 1), to unambiguously identify the intrinsically bound metal ions in the active site of the PTE. The data showed unequivocally that the two metal ions within the active site are indeed Zn atoms, confirming their expected identity (Figure 12).

**Figure 11.**
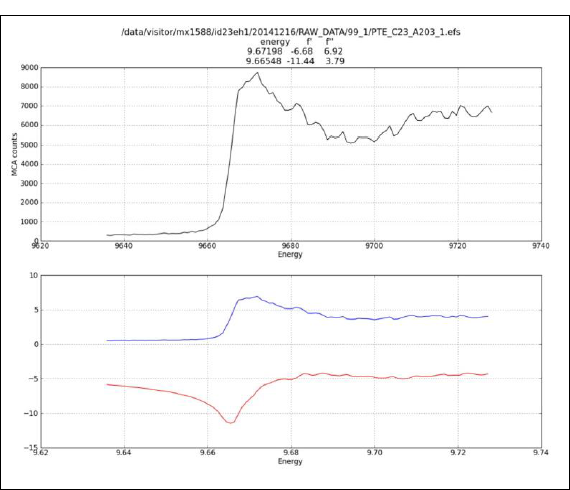
Scan of the C23M_1 crystal at a range of energy showing a peak corresponding to the Zn absorption edge (top). Scattering factors (f’ and f’’) as a function of energy (bottom).

**Figure 12.**
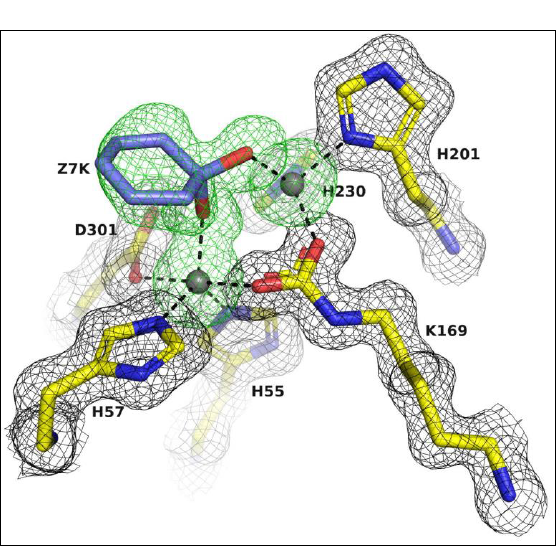
Electron density omit maps of the active-site region of the C23M_1 structure exhibit. The two Zn^+2^ ions and the unknown blob of electron density labeled 7ZK were omitted in calculating the electron densities. The 2Fo-Fc omit map, contoured at 1 σ, is shown in grey. The Fo-Fc omit map, contoured at 2.5 σ, is shown in green.

### 3.6. A cyclic compound in the PTE active site

It is not uncommon for crystallization reagents to be observed in the active site of protein structures. In the case of the PTE variants, PEG and MPD were used as precipitants for crystallization A53_1, A53_3, C23_1 (Table 1), and C23M (data not shown). As described above A53_1 is presumably a nonactive monomer with one Zn^+2^ atom in the active site. C23_1 diffracted only to 3.2 Å resolution and A53_3 crystallized in the *P2_1_* space group with the residual octapeptide in one subunit pointing into the active site of a subunit symmetry related. The A53_3 apo structure also has an unidentified six-member ring ligand (D9K) in its active site (Figure 12). D9K has two oxygen atoms likely acting as a chelator that form coordinate bonds with the two Zn^+2^ ions in the solution and may have been carried into the crystals during protein expression and purification. Another unknown six-member ring ligand is observed in the apo structures of A53 and C23M crystallized from AS and glycerol, A53_2 (Q5K) and C23M_1 (Z7K) (Table 1). Similar to D9K also the Q5K and Z7K acts as a chelator by creating coordinate bonds with Zn^+2^ ions. Notably, no OP ligands were observed in the active site of the variant crystallized from PEG with MPD (data not shown).

### 3.7. Co-crystallization and soaking of OPs to PTEs

Soaking crystals with ligands is a common method to obtain the structure of protein-ligand complexes, but it requires careful consideration of several factors, including the requirement to dissolve the ligand in the crystallization reagents or in a solvent that won’t destroy the protein crystal, soaking time and the ligand concentration. One significant limitation of soaking is the requirement for a crystal form with an accessible ligand-binding site or a bound ligand that can be easily exchanged with a different ligand. In the case of the PTE variants, attempts to soak OPs into existing crystals of the native enzymes were unsuccessful because the active site was already occupied by ligands (*i.e*. PAA or unidentified six-member ring ligand shown in Figures 10 and in 12 and 13 respectively) coordinated to the α-Zn^2+^ and β-Zn^2+^ ions, making it difficult for OPs to replace them. Furthermore, the presence of a flexible residual octapeptide linker close to the active site (Figure 9) could have also interfered with the ligand-binding site, further complicating the soaking experiments.

**Figure 13.**
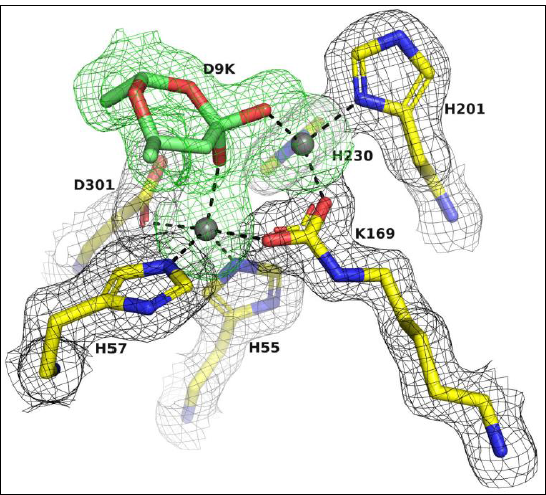
Figure 13 Electron density omit maps of the active-site region of the A53_3 (Chain A) structure exhibit. The two Zn^+2^ ions and the unknown blob of electron density labeled D9K were omitted in the calculation of the electron densities. The 2Fo-Fc omit map, contoured at 1 σ, is shown in grey. The Fo-Fc omit map, contoured at 2.5 σ, shown in green.

Failure of the OPs to displace the ligands in the active site could be due either to the lack of flexibility of the crystalline protein or to crystal-packing constraints. Also, when performing ‘replacement soaking,’ one has to consider and compare the binding affinity of the soaked ligand to that of the OPs to the PTEs. In the case where the flexible residual octapeptide linker was observed close to the active site, it possibly interfered with the ligand-binding site explaining why no OPs were observed.

Co-crystallization, on the other hand, typically requires more resources in terms of time and ligand and protein consumption. However, when the ligands are insoluble or the protein aggregates easily, co-crystallization is the method of choice. Co-crystallization is also preferred when ligand binding is associated with conformational changes, and the active site is occupied with a ligand that cannot be displaced by another ligand. In the PTE variants studied, OPs could only be observed in the active site when co-crystallization experiments were performed, and only in crystals obtained from AS with glycerol. The electron density in the active site of the A53_5, C23_3, and the C23M_2 (Figure 14) structures co-crystallized with methylphosphonate clearly indicated that the ligand was replacing the six-member ring compound present in the apo structures (A53_2, and C23M_1). Similarly, the O-ethyl methylphosphonate was observed in the active site of the C23_5 structure, the C23_4 active site shows electron density corresponding to part of the O-ethyl-O-(N,N-diisopropylaminoethyl) methylphosphonate and part of the O-isopropyl methylphosphonate was observed in C23_2.

**Figure 14.**
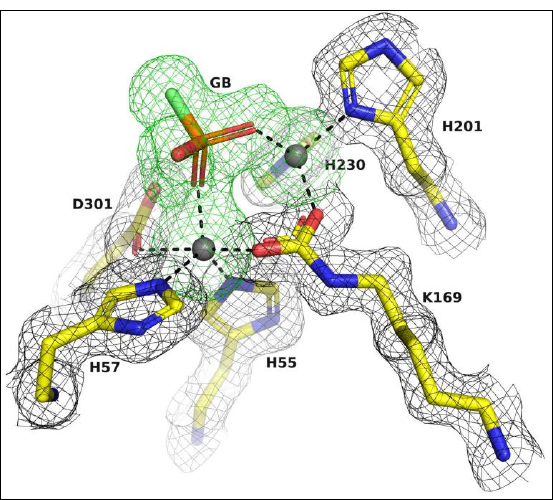
Electron density omit maps of the active-site region of the C23M_2 structure exhibit. The two Zn^+2^ ions and the methylphosphonate labeled GB were omitted in the calculation of the electron densities. The 2Fo-Fc omit map, contoured at 1 σ, is shown in grey. The Fo-Fc omit map, contoured at 2.5 σ, is shown in green.

## 4. Discussion

Our study describes the challenges encountered in obtaining crystal structures of the enzyme PTE in complex with organophosphate (OP) ligands. Crystallization solutions contain a spectrum of chemicals that act as protein precipitants, buffers, and/or reagents to increase protein stability. The effective metal concentration in the solution may be decreased by the formation of metal complexes with these chemicals. Many transition metal ions will form insoluble hydroxides in neutral to basic pH, which can significantly limit metal solubility and substantially decrease the effective metal concentration. Besides the formation of insoluble hydroxides in solution, some crystallization reagents can act as chelators by creating coordinate bonds with metal ions in solution. Therefore, it is essential to carefully consider the choice and concentration of reagents used in the crystallization process to ensure that they do not interfere with the function of the target protein.

Zinc is required for the activity of > 300 enzymes, covering all six classes of enzymes (McCall *et al*., 2000), one of which is PTE. We observed that the A53_1 variant was a non-active monomer with only one Zn^2+^ ion in the active site. To counteract this problem, it was necessary to increase the concentration of ZnCl_2_ in the purification and crystallization steps. The fact that the presence of the two Zn^+2^ ions was crucial for the formation of the canonical PTE dimer structure and for its activity implies that the Zinc ions participate directly in chemical catalysis as well as for maintaining protein structure and its stability. The identity of the Zn^2+^ ion in the PTE active site was confirmed by collecting X-ray data at the Zn absorption edge.

This study has shown that some of the crystallization precipitants and compounds used in the protein expression and purification processes could lodge within the PTE active site, competing with and even precluding ligand binding. Specifically, the PAA used in the crystallization solution was detected in the active site of the PTE variants and a well-defined electron density of an unidentified six-member ring compound was observed in the PTE active site when either PEG 6000 with MPD or AS with glycerol were employed in the crystallization trials. Soaking OPs to the crystals of the apo PTE variants grown from the three different crystallization conditions failed to replace either the 6-member ring or the PAA observed in the active site. Furthermore, co-crystallization experiments of the OP ligands with the A53, C23, or C23M variants produce crystals grown from the same conditions as those of the Apo. To our surprise also, in the co-crystallization experiments, the PAA and the 6-member ring lodge within the PTE active site and impeded OP binding. This might suggest that the PTEs had a higher affinity to the crystallization compounds than to the OPs. Only when the PTEs and the OPs are co-crystallized from AS with glycerol that the OPs are observed in the active site. The electron density for the methylphosphonate was observed in the active site of A53_5, C23_3, and C23M_2 structures, while the O-ethyl methylphosphonate was observed in the active site of C23_5 structure. Furthermore, the C23_4 active site showed electron density corresponding to part of the O-ethyl-O-(N,N-diisopropylaminoethyl) methylphosphonate and part of the O-isopropyl methylphosphonate was observed in C23_2. Together these observations indicate that the choice of crystallization conditions was crucial for obtaining OP complexes.

To increase PTE expression levels and solubility, maltose-binding protein (MBP) was introduced as a fusion partner and was removed by digestion with Factor Xa prior to crystallization trials. Unexpectedly, electron density corresponding to the residual octapeptide spacer penetrated the PTE active site, thereby blocking entry to the active site thus preventing the binding of OPs. Interestingly, the residual octapeptide was observed in PTE crystals obtained from different crystallization precipitates but all crystallized in the same space group (P2_1_) implying that the space group in which a protein crystallizes can affect the conformation of the protein and the binding of ligands. This is because the space group determines the crystal packing, which in turn affects the local environment around the protein molecules. Our findings illustrate that the space group is a pivotal factor in positioning the residual octapeptide in the PTE active site and thereby impending OP binding. However, it is noteworthy that the space group by which a protein is crystallized is not typically taken into account in computer-aided drug design methods used to identify promising drug candidates, which rely on the protein structure alone to predict binding. Furthermore, our results also highlight the importance of removing residual tags used to increase expression and purification levels before attempting to study the 3D structure of proteins with bound ligands. The presence of these tags can penetrate the active site and hinder ligand binding, as demonstrated in the case of the PTE.

Our findings highlight the crucial role of various factors such as the molecular constructs, crystallization conditions, space groups, compounds carried from protein expression and purification, and the presence of residual tags in accurately visualizing molecular complexes of ligands bound to the enzyme. The findings of the study also underscore the importance of careful experimental design and rigorous data analysis to ensure the accuracy and reliability of resulting protein-ligand structures. The study demonstrates that the presence of crystallization precipitants can compete with and even preclude ligand binding, leading to incorrect identification of lead drug candidates. Therefore, it is essential to consider various factors and optimize experimental conditions to obtain accurate and reliable results in the study of protein-ligand interactions and drug design.

## Acknowledgements

This work benefited from access to the Structural Proteomics Unit at the Weizmann Institute of Science, an Instruct-ERIC center. Financial support was provided by the Center for ScientificExcellence at the Weizmann Institute of Science.

## References

Afonine, P. V., Grosse-Kunstleve, R. W., Echols, N., Headd, J. J., Moriarty, N. W., Mustyakimov, M., Terwilliger, T. C., Urzhumtsev, A., Zwart, P. H. & Adams, P. D. (2012). Acta Crystallogr D Biol Crystallogr 68, 352–67.

Arnold, F. H. & Haymore, B. L. (1991). Science 252, 1796–1797.

Benning, M. M., Kuo, J. M., Raushel, F. M. & Holden, H. M. (1994). Biochemistry 33, 15001–15007.

Benning, M. M., Shim, H., Raushel, F. M. & Holden, H. M. (2001). Biochemistry 40, 2712-2722.

Berman, H. M., Westbrook, J., Feng, Z., Gilliland, G., Bhat, T. N., Weissig, H., Shindyalov, I. N. & Bourne, P. E. (2000). Nucleic Acids Res. 28, 235–242.

Bigley, A. N., Mabanglo, M. F., Harvey, S. P. & Raushel, F. M. (2015). Biochemistry 54, 5502–5512.

Bigley, A. N., Xu, C., Henderson, T. J., Harvey, S. P. & Raushel, F. M. (2013). J. Am. Chem. Soc. 135, 10426–10432.

Chen, V. B., Arendall, W. B., 3rd, Headd, J. J., Keedy, D. A., Immormino, R. M., Kapral, G. J., Murray, L. W., Richardson, J. S. & Richardson, D. C. (2010). Acta Crystallogr. D Biol. Crystallogr. 66, 12-21.

Cherny, I., Greisen, P., Jr., Ashani, Y., Khare, S. D., Oberdorfer, G., Leader, H., Baker, D. & Tawfik, D. S. (2013). ACS Chem. Biol. 8, 2394–2403.

Corpet, F. (1988). Nucleic Acids Res. 16, 10881–10890.

Dai, J., Bao, Z., Sun, L., Hong, S. U., Baker, G. L. & Bruening, M. L. (2006). Langmuir 22, 4274–4281.

Dym, O., Song, W., Felder, C., Roth, E., Shnyrov, V., Ashani, Y., Xu, Y., Joosten, R. P., Weiner, L., Sussman, J. L. & Silman, I. (2016). Protein Sci. 25, 1096–1114.

Emsley, P. & Cowtan, K. (2004). Acta Crystallogr. D Biol. Crystallogr. 60, 2126–2132.

Evans, P. (2006). Acta Crystallogr. D Biol. Crystallogr. 62, 72-82.

Fischer, B. E., Haring, U. K., Tribolet, R. & Sigel, H. (1979). Eur. J. Biochem. 94, 523–530.

French, S. & Wilson, K. (1978). Acta Crystallographica Section A: Foundations and Advances 34, 517–525.

Goldsmith, M., Aggarwal, N., Ashani, Y., Jubran, H., Greisen, P. J., Ovchinnikov, S., Leader, H., Baker, D., Sussman, J. L., Goldenzweig, A., Fleishman, S. J. & Tawfik, D. S. (2017). Protein Eng. Des. Sel. 30, 333–345.

Goldsmith, M., Eckstein, S., Ashani, Y., Greisen, P., Jr., Leader, H., Sussman, J. L., Aggarwal, N., Ovchinnikov, S., Tawfik, D. S., Baker, D., Thiermann, H. & Worek, F. (2016). Arch. Toxicol. 90, 2711–2724.

Grimsley, J. K., Calamini, B., Wild, J. R. & Mesecar, A. D. (2005). Arch. Biochem. Biophys. 442, 169–179.

Harper, L. L., McDaniel, C. S., Miller, C. E. & Wild, J. R. (1988). Appl. Environ. Microbiol. 54, 2586–2589.

Holm, L. & Sander, C. (1997). Proteins 28, 72–82.

Horne, I., Sutherland, T. D., Harcourt, R. L., Russell, R. J. & Oakeshott, J. G. (2002). Appl. Environ. Microbiol. 68, 3371–3376.

Jackson, C. J., Weir, K., Herlt, A., Khurana, J., Sutherland, T. D., Horne, I., Easton, C., Russell, R. J., Scott, C. & Oakeshott, J. G. (2009). Applied and Environmental Microbiology 75, 5153–5156.

Joosten, R. P., Long, F., Murshudov, G. N. & Perrakis, A. (2014). IUCrJ 1, 213–220.

Leslie, A. & Powell, H. (2007). in Evolving Methods for Macromolecular Crystallography, (R. J. Read & J. Sussman, Eds), Springer Netherlands: Dordrecht. pp. 41-51.

Masson, P. & Rochu, D. (2009). in Handbook of Toxicology of Chemical Warfare Agents, (R. C. Gupta, Eds), Academic Press: San Diego. pp. 1053-1065.

McCall, K. A., Huang, C.-c. & Fierke, C. A. (2000). J. Nutr. 130, 1437S-1446S.

McCoy, A. J., Grosse-Kunstleve, R. W., Adams, P. D., Winn, M. D., Storoni, L. C. & Read, R. J. (2007). J. Appl. Cryst. 40, 658–674.

McLoughlin, S. Y., Jackson, C., Liu, J. W. & Ollis, D. (2005). Protein Expr. Purif. 41, 433–440.

Murshudov, G. N., Vagin, A. A., Lebedev, A., Wilson, K. S. & Dodson, E. J. (1999). Acta Crystallogr. D Biol. Crystallogr. D55, 247–255.

Robert, X. & Gouet, P. (2014). Nucleic Acids Res. 42, W320–324.

Singh, B. K. (2009). Nature reviews. Microbiology 7, 156-164.

Sussman, J. L., Lin, D., Jiang, J., Manning, N. O., Prilusky, J., Ritter, O. & Abola, E. E. (1998). Acta Crystallogr. D Biol. Crystallogr. 54, 1078–1084.

Tokuriki, N., Jackson, C. J., Afriat-Jurnou, L., Wyganowski, K. T., Tang, R. & Tawfik, D. S. (2012). Nat. Commun. 3, 1257.

Tsai, P. C., Fox, N., Bigley, A. N., Harvey, S. P., Barondeau, D. P. & Raushel, F. M. (2012). Biochemistry 51, 6463–6475.

Yang, C. Y., Renfrew, P. D., Olsen, A. J., Zhang, M., Yuvienco, C., Bonneau, R. & Montclare, J. K. (2014). ChemBioChem 15, 1761–1764.

Yang, H., Carr, P. D., McLoughlin, S. Y., Liu, J. W., Horne, I., Qiu, X., Jeffries, C. M., Russell, R. J., Oakeshott, J. G. & Ollis, D. L. (2003). Protein Eng. 16, 135–145.

Zhao, X., Li, G. & Liang, S. (2013). Journal of Analytical Methods in Chemistry 2013, 581093.

Zheng, H., Chruszcz, M., Lasota, P., Lebioda, L. & Minor, W. (2008). J. Inorg. Biochem. 102, 1765–1776.

